# Ordered release of genomic RNA during icosahedral virus disassembly

**DOI:** 10.1101/2025.01.09.632012

**Authors:** Yiyang Zhou, Andrew L. Routh

**Affiliations:** Department of Microbiology and Immunology, The University of Texas Medical Branch, Galveston, Texas, USA; Department of Biochemistry and Molecular Biology, The University of Texas Medical Branch, Galveston, Texas, USA; Department of Immunology and Microbiology, Scripps Research, La Jolla, California, USA; Sealy Center for Structural Biology and Molecular Biophysics, The University of Texas Medical Branch, Galveston, Texas, USA; Institute for Human Infections and Immunity, University of Texas Medical Branch, Galveston, Texas, USA

**Keywords:** icosahedral virus, RNA release, virus disassembly, PT-ClickSeq, RNA-capsid interaction

## Abstract

To release their genomic cargo, many icosahedral viruses undergo a series of ordered conformational changes via distinct disassembly intermediates that allow nucleic acid egress. Previous studies have focused on the rearrangement of the virus capsid protein shell during disassembly. However, it is unclear whether the packaged viral nucleic acids also undergo defined rearrangements or whether specific regions of the viral genome are released in a predetermined order. In this study, we established a Next-Generation Sequencing platform (“*PT-ClickSeq*”) that does not require RNA extraction or fragmentation and so can natively sequence the nucleic acid exposed during virus particle disassembly without disrupting the capsid protein. We used Flock House virus (FHV) as a model system, which produces two well-defined disassembly intermediates with externalized RNA. With PT-ClickSeq, we found that FHV releases RNA genome in an ordered and conserved fashion with the 5’ and 3’ terminal regions of viral genomic RNAs and specific loci within the viral genome being released first during virus particle disassembly. Further, different genomic loci exhibit distinct energy barriers prior to release, suggesting a programmed exposure of the viral RNA. We also characterized viral RNA-capsid interactions using established cross-linking and NGS approaches (“vPAR-CL”) and observed a disordering of RNA-capsid interactions during disassembly. Interestingly, we noticed an anti-correlation between preferentially released RNA regions and strong RNA-capsid interactions observed prior to disassembly. These findings demonstrate that rather than being a passive cargo, the encapsidated viral genome serves an important role in programmed virus disassembly.

**Significance:** Viruses need to strike a balance between structural rigidity and flexibility, to achieve both sufficient protection and rapid release of packaged genome into host cells. During the process of genome delivery, many viruses undergo a programed disassembly process through successive morphological changes, which give rise to partially dissembled virus particles, termed disassembly intermediates. It is important to study these intermediates as “checkpoints” to understand virus disassembly dynamics. We established a next-generation sequencing method that can monitor the RNA behavior during these conformational changes. We found that different regions of RNA were released with different energy thresholds and the RNA release prioritized regions with low RNA-protein interactions. These findings shed light on the active role of the viral RNA in virus disassembly.

## Introduction

An interesting duality for non-enveloped icosahedral viruses is that the capsid shell is required to provide structural stability and protection of the encapsidated genetic cargo to external environmental factors to allow dissemination between hosts, but must also allow rapid release of the genomic cargo into a host cell at the appropriate time during an infection. For non-enveloped viruses, switching between these two distinct roles is critical to avert premature exposure of the genomic cargo and to ensure its delivery to the correct cellular compartment [1]. To achieve this, viruses employ diverse mechanisms that trigger virus particle disassembly and genome release, such as receptor binding [2–5], compartmental pH changes [6–9] and other mechanical cues [1, 10, 11].

During the disassembly process, some viruses may proceed via an “*en masse*” approach, characterized by a heterogeneous and rapid disintegration of the capsid shell into component subunits [12, 13]. Conversely, viruses may disassemble via a coordinated step-wise process, comprising successive conformational changes through multiple intermediate particles. This is demonstrated by polioviruses, rhinoviruses and other enteroviruses, where transition from a full 160S virion to a 135S particle (or “A particle”) is triggered by receptor binding in physiological temperature during early infection cycle. The “A particle” is further deconstructed into an 80S particle after RNA genome release [2]. Similarly, multi-layered viruses, such as reoviruses and rotaviruses, sequentially lose capsid layers after receptor binding or during membrane penetration [14, 15]. Single-layered icosahedral viruses may also shed specific capsid subunits (such as monomers, pentamers, hexamers or others) to give rise to specific structural intermediates that feature a rupture or pore in the capsid shell through which the genomic cargo may egress.

Disassembly intermediates provide insightful timestamps with which to characterize the spatiotemporal disassembly process of non-enveloped viruses. However, while previous studies have focused on the capsid proteins and their higher-order structures during disassembly, little is known about whether the genomic material is released in an ordered or structurally conserved manner. The transient and heterogeneous nature of intermediate particles presents a significant challenge in structural approaches such as crystallography or cryoEM. Previous studies have achieved high-resolution asymmetrical reconstructions of disassembly intermediates and revealed conformational changes in the genetic cargo inside the viral capsid[16]. However, these studies do not determine whether different genomic region might be preferentially released during particle disassembly, or the sequence identity of genomic RNA responsible for the observed RNA conformations.

Flock House virus (FHV, *Nodaviridae*) provides a model icosahedral virus for understanding non-enveloped virus assembly, disassembly and RNA-capsid dynamics [17–20] and has well-documented disassembly intermediates in controlled experimental settings [16]. The *T* = 3 icosahedral FHV particle stoichiometrically packages one molecule of each of the two genomic segments (RNA1 of 3.1 kb and RNA2 of 1.4kb), which collectively encodes only 3 viral proteins: (RNA dependent RNA polymerase and protein B2 on RNA1, capsid on RNA2). Despite the simple particle structure and small genome, it has been demonstrated that FHV virion is packed with extensive RNA-protein interactions [20]. A recent study induced FHV particle rearrangements with incremental heating *in vitro*, and characterized with cryoEM the transformation from intact virions into “eluted” and a further “puff” disassembly intermediates [16]. Interestingly, FHV “puff” intermediate particles showed further disintegration of capsid layer and a partial RNA genome release through a 2-fold axis (hence the name “puff”).

In this study, we continue to use Flock House virus (FHV) as a model system to study the molecular mechanisms and events associated with RNA virus entry and particle disassembly. We repurposed a Next-Generation Sequencing (NGS) technique ‘ClickSeq’, which is characterized by its use of “click-chemistry” for sequencing adaptor ligation. An important feature of ClickSeq is the avoidance of pre-treatment or fragmentation of nucleic acids prior to library synthesis [21]. This allows NGS library preparation directly from unextracted biological materials, which we term Particle-Templated ClickSeq (“PT-ClickSeq”). We demonstrate here that PT-ClickSeq is a simple and qualitative method that reports on which regions of RNA are exposed in an ensemble of RNA-complexes, such as viral disassembly intermediates.

With PT-ClickSeq, we found that the specific regions of the FHV genomic RNA are exposed to different extents in FHV disassembly intermediate particles. We demonstrated that the release of FHV RNA genome followed a step-wise pattern and that different genomic regions required different energy thresholds to be released. We further utilized viral Photo-Activatable Ribonucleoside CrossLinking (vPAR-CL) [20, 22] to characterize the RNA-capsid interactions during FHV disassembly. We found that the preferentially exposed RNA loci lack significant RNA-capsid interactions. This indicates an active role for the viral genome in directing the choreographed structural details of virus particle disassembly.

## Results

### FHV disassembly intermediate particles are generated *in vitro* via heat-treatment

The formation of specific FHV disassembly intermediate particles can be induced upon controlled heating of wild-type virions *in vitro*, with two successive conformational transitions at 70 °C and 75°C. These distinct structural states have previously been characterized by negative-stain transmission electron microscopy and cryo-electron microscopy with three-dimensional image reconstruction [16]. Schematically (**Fig. 1a**): wild type (wt) FHV comprises an ultra-stable [23], non-enveloped virion approximately 34 nm in diameter with extensive RNA-capsid interactions between the well-defined T=3 icosahedral capsid shell and dodecahedral cage of encapsidated RNA [17, 20, 24, 25]. When FHV virions are heated to 70 °C, particles show a reduction in both density and dimension, and particles become permeable to negative stain, forming the “eluted” particles[16]. When FHV virions are heated to 75 °C, the capsid shell continues to lose structural integrity and a portion of encapsidated RNA genome can be observed extruding from the capsid shell to form “puff” particles[16].

**Figure 1:**
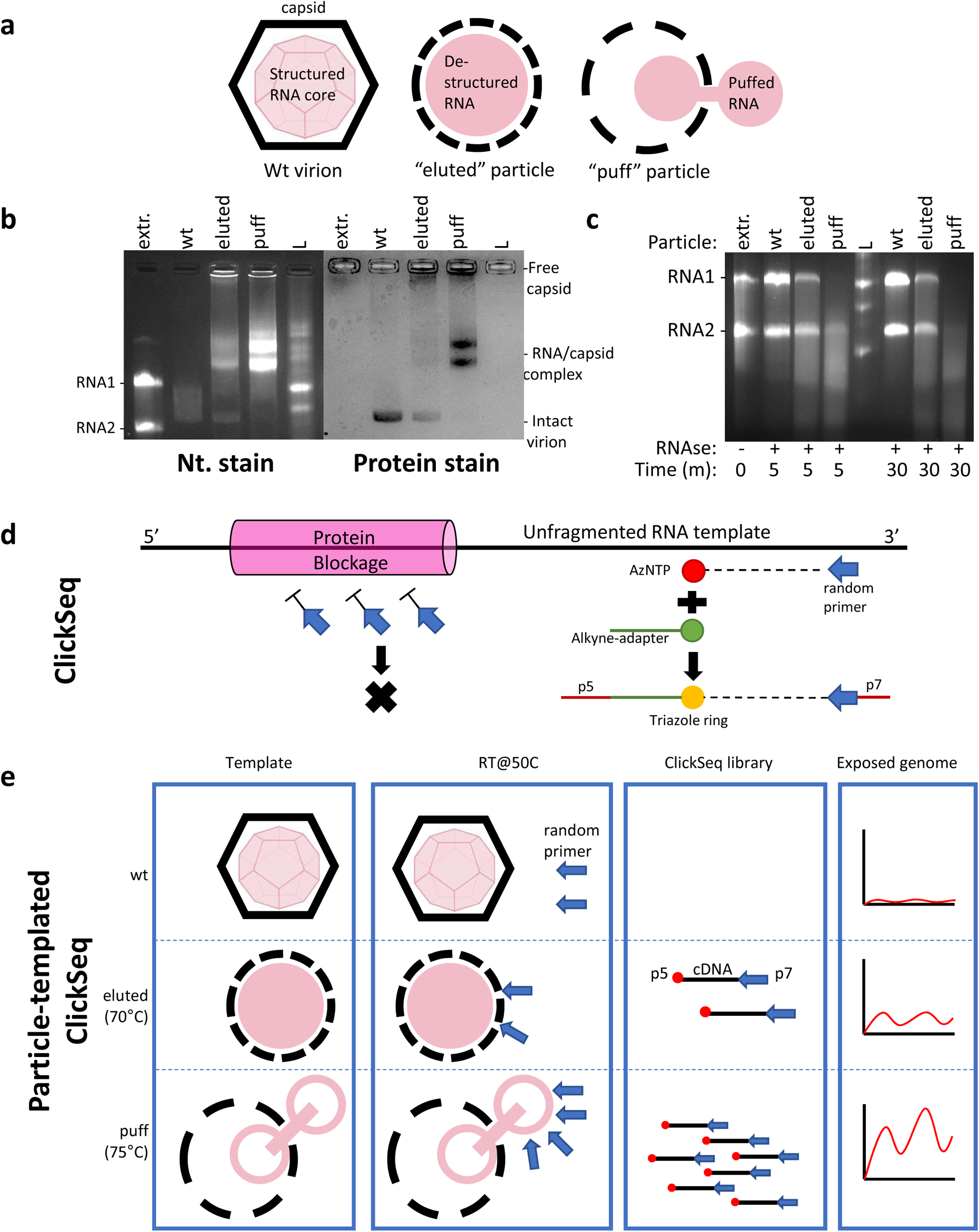
Sequencing of RNA exposed in disassembly intermediates of virus particles. (**a**) Schematic of Flock House virus (FHV) wildtype (wt), ‘*eluted’* and ‘*puff’* particles. (**b**) Different particles show different permissiveness to nucleic acid stain in a non-denaturing RNA electrophoretic gel. Both eluted and puff particles form multiple RNA-capsid complexes, visualized using both nucleic acid stain (GelStar, left) and protein stain (Coomassie, right). (**c**) The RNA genome of different particles showed different susceptibility to RNase A digestion. “Extr” = purified RNA extracted from wt particles. “L” = ssRNA ladder. (**d**) Schematic of ClickSeq approach to assay exposed RNA. (**e**) Particle-templated ClickSeq (PT-ClickSeq) uses RNA-protein complexes (such as a FHV puff particle) directly as the template during RT reaction to probe and sequence the exposed RNA regions. The different levels of RNA genome exposure are reflected by disproportionate sequence coverages. “p5” and “p7” = Illumina sequencing adapter(s)

In this study, we recreated the FHV eluted and puff intermediate particles with controlled incremental heating methods as described previously [16]. 50 µg of wt, eluted and puff particles were directly loaded onto a 1% non-denaturing agarose gel pre-stained with nucleic acid dye (**Fig. 1b**, left). Intact wt virions exhibited little permeability to nucleic acid stain and appeared as a faint smear. This is consistent with previous thermostability studies of FHV [26]. In contrast, eluted particles consisted of a mixture of intact wt virions and heterogeneous RNA-capsid complexes, which migrated slower than extracted and purified genomic RNA. The heterogeneous population of the eluted particles is consistent with a previous characterization of negative-stained low-resolution micrographs [16], although in our molecular assay, the percentage of wt virions was higher. In our assay, puff particles showed evidence of a continued disintegration from eluted particles. This is reflected by the increased nucleic acid stain of the RNA-capsid complex(es), altered migration pattern, and a lack of evidence of wt virions. Next, the same gel was post-stained with Coomassie Blue to evaluate the protein content of each sample (**Fig. 1b**, right). Puff particles exhibited stronger Coomassie staining than the eluted particles, potentially due to the increased disorganization of capsids in higher temperature. FHV is known to embrace extensive RNA-capsid interactions in wt virion [20]. It is striking that, in spite of excessive heating (70°C/75°C for 30 min, respectively), eluted/puff particles retained the association of capsid protein with the viral RNA.

### FHV disassembly intermediate particles are structurally stable

In wt FHV, the intact icosahedral shell protects the encapsidated RNA from environmental factors such as naturally occurring RNases that would lead to genomic RNA digestion or fragmentation. We sought to determine whether eluted or puff particles could retain such RNase resistance. 50 µg of purified virus particles were treated with 5µg of RNase A for at 37 °C, and the remaining viral RNAs were extracted thereafter and analyzed by electrophoresis on a non-denaturing agarose gel (**Fig. 1c**). In spite of RNase A treatment, eluted particles retained some undigested RNA1 and RNA2, albeit at yields less than that of wt virion. This is consistent with a partial protection of genomic RNA. In addition, eluted particles also showed a substantially greater amount of RNA smearing at lower molecular weights. This is consistent with above-stated observation that eluted particles are structurally heterogeneous. In contrast, Puff particles retained only a small amount of full length RNA2 after RNase digestion, with a higher intensity of lower molecular weight RNA smears relative to that of the eluted particles. This indicates that in puff particles, RNA1 is more susceptible to RNase digestion than RNA2. We further extended the RNase digestion time to 30 min, which showed similar results to 5 min digestion. This suggests that eluted and puff particles were relatively stable during this extended treatment and only part of their genomes were susceptible to RNase digestion and that digestion of the exposed RNA does not trigger further disassembly of the virus particles. These results demonstrate that FHV disassembly intermediates are structurally stable, and exhibit altered RNA-capsid interactions.

### Particle-Templated ClickSeq (PT-ClickSeq) sequences exposed RNA in intermediate particles

ClickSeq is a Next-Generation Sequencing (NGS) library preparation method characterized by the use of click-chemistry in place of enzymatic ligation for the addition of sequencing adaptors [21, 27]. Unlike conventional NGS library preparations, ClickSeq does not require fragmentation or any other pre-treatment of the input RNA template. In ClickSeq, a randomly-primed RT reaction is supplemented with 3’-azido-dNTPs that randomly terminate cDNA synthesis and release 3’-azido blocked cDNA fragments. Such 3’-azido-cDNAs are then “click-ligated” to a 5’-alkyne-functionalized sequencing adapter via click chemistry (Copper-catalyzed Azide-Alkyne Cycloaddition, CuAAC). After final PCR of the “click-ligated” cDNA, this process generates an NGS library with the required (e.g. Illumina) sequencing adaptors and indexes (**Fig. 1d**). As ClickSeq does not require pre-treatment of the RNA or any RNA extraction prior to reverse transcription, ClickSeq can utilize a partially dissembled virus particle as template (**Fig. 1e**). This averts denaturing the RNA-protein complex during RNA extraction and retains the particle structure of disassembly intermediates. During RT, the exposed RNAs in disassembly intermediates allow the annealing of random hexamers, while the protected RNAs impede primer annealing. As a result, only exposed, unprotected RNA can be successfully converted into an cDNA library and subsequently sequenced. Therefore, the uneven genomic coverage of PT-ClickSeq can reveal which regions of the genomic RNA are exposed in the different disassembly intermediates of FHV.

### PT-ClickSeq reveals differential release of genomic RNA in disassembly intermediates

We first established the baseline of PT-ClickSeq read coverage over unencumbered, protein-free RNA. For this, purified RNAs extracted from wt virions (n=3) were used. The mapped reads were ratiometrically normalized to the total sequence depth. Consistent with prior NGS studies of FHV genomic RNA[28], the extracted FHV RNA demonstrated even read depth mapping across the viral genome (**Fig. 2a, f**, green dashed line), with expected minor undulations in read coverage indicative of potential biases inherent to NGS approaches due to viral primary sequence (e.g. GC content), in uneven primer annealing, and/or RNA secondary structures.

**Figure 2:**
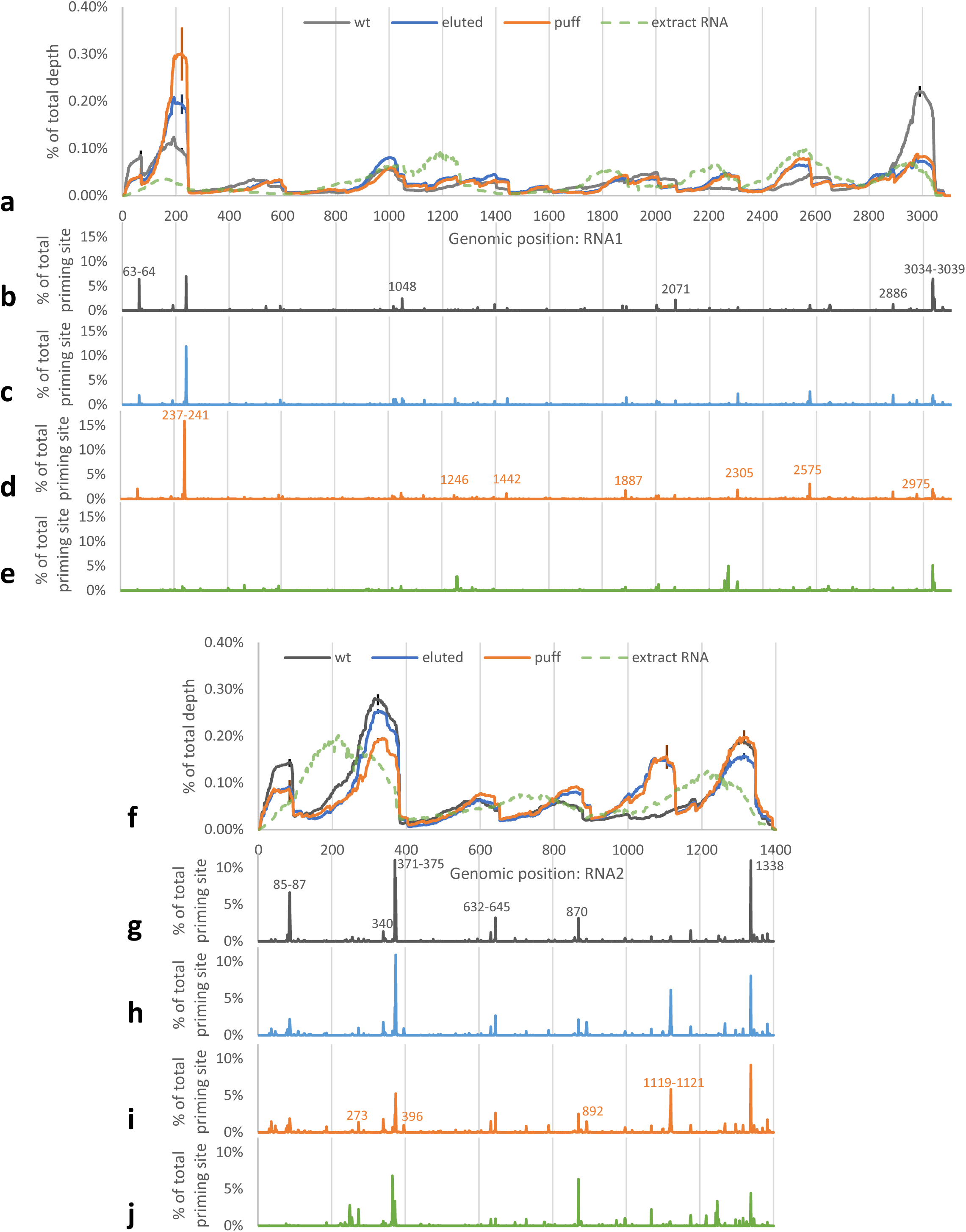
Particle-templated ClickSeq (PT-ClickSeq) reveals different genome coverages for disassembly intermediates. Wt, eluted and puff particles show distinct unevenness in read coverage for both RNA1 (**a**) and RNA2 (**f**). In contrast, extracted and purified genomic RNA is characterized by even and smoother read coverage across genomes. With paired-end sequencing, the percentage of priming sites on RNA1 (**b-e**) and RNA2 (**g-j**) is also consistent with overall genomic coverage. (Error bars: standard deviation, N=3, only displayed on representative positions for clarity)

After establishing a baseline, we sought to determine which regions of the FHV genomic RNAs were preferentially exposed in FHV disassembly intermediates. FHV wt (no-preheating), eluted (preheated at 70°C) and puff particles (preheated at 75°C) were directly used as input without RNA extraction in PT-ClickSeq (n=3 for each condition). Libraries were successfully generated for each sample, although a lower yield of final NGS library for the FHV wt sample likely reflected the reduced availability of viral genomic RNA.

In contrast to the extracted FHV RNAs, sequencing of the viral disassembly intermediate samples revealed a stark unevenness in the distribution of sequence reads across the viral genomic RNAs, indicative of different genome exposure among wt, eluted and puff particles for both RNA1 and RNA2 (**Fig. 2a, f**). For RNA1, the sequence read coverage for the wt particles showed clear enrichment at both 5’- and 3’-terminal regions, while the rest of genome yielded relatively little read depth (<0.1% of total depth at each given nucleotide position). More specifically: RNA1: 26-75 nts (1.6% of RNA1 length) collectively accounted for 3.6% of total RNA1 coverage; RNA1: 151-250 nts (3.2% of RNA1 length) collectively accounted for 9.7% of total RNA1 coverage; RNA1: 2901-3050 nts (4.8% of RNA1 length) collectively accounted for 23.6% of total RNA1 coverage (**Supplemental Fig. 1**). In contrast, the read coverage of extracted and purified genomic RNA was relatively even in these same regions (**Supplemental Fig. 1**). As the coverage information was normalized to the percentage of total sequenced depth, this suggests that the exposed RNAs were disproportionally localized at the 5’ and 3’ proximities of wt particles but not at the rest of genome.

Similarly to wt FHV, both eluted and puff particles showed increased coverage at the 5’ terminal 26-75nts (1.8% of total coverage) and 3’-terminal 2901-3050nts (9.0-9.7% of total coverage) of RNA1 relative to the ‘naked’ FHV RNA (**Supplemental Fig. 1**). The most dramatic increase of read coverage was observed at 151-250 nts of RNA1, accounting for 16.8 and 23.3% of the total mapped reads for eluted and puff particles respectively, while only 2.7% of the reads mapped in this region for the ‘naked’ FHV RNA (**Supplemental Fig. 1**). In addition, we also detected increases of read coverage at 2200-2300 nts and 2425-2725 nts in both eluted and puff particles.

In RNA2, wt FHV particles again showed disproportionally high coverage at 5’ terminal 26-100nts (5.4% of RNA2 length and 8.9% of total RNA2 coverage), 276-375nts (7.1% of RNA2 length and 24.9% of total RNA2 coverage) and 3’ terminus 1251-1350nts (7.1% of RNA2 length and 16.2% of total RNA2 coverage) (**Supplemental Fig. 1**). In contrast, eluted and puff particles exhibited a substantial increase in coverage in RNA2: 1051-1150nts. (wt: 3.6%, eluted: 11.8%, puff: 11.7%), and a slight increase in 800-875nts.

Importantly, the disproportionate enrichment of sequence data over these regions is not seen in the extracted FHV RNA samples and exceeds the magnitude of the ‘undulations’ of sequence coverage in the baseline. This provides confidence that these enrichments are due to the increased exposure of released RNA and de-encapsidation of RNA in the disassembly intermediates, rather than due to artefacts and biases of NGS library prep and PCR. To further demonstrate the above-stated observations, a moving average map (50 nts. window) of genomic coverage and the collectively coverage of specific genomic regions is provided in **Supplemental Fig. S1b.**

### PT-ClickSeq identifies priming sites of exposed RNA

As a complementary analysis to scrutinizing read coverage, we further investigated the paired-end read sequences of PT-ClickSeq data. The 5’-most coordinates of the ‘R2’ read of the paired-end sequences revealed the priming sites during the reverse transcription reaction of different disassembly intermediates (**Supplemental Figure S2**, priming site is defined here as the immediate upstream site of 3’ random hexamer sequence). The normalized priming site depths (**Fig. 2b-e, g-j**) showed patterns consistent with the genomic coverage data (**Fig. 2a, f**). Several abundant priming sites were located directly downstream of high genomic coverage regions and their depths ratiometrically corresponded to the coverage of these regions. These paired-end data further demonstrate that the coverage differences in wt, eluted, and puff particles were led by the different RNA exposure levels and resultant primer annealing frequencies.

The coverage and priming site discrepancies revealed by PT-ClickSeq provide several insights into FHV disassembly intermediates and genome release: (**A**) the differential genome release at low temperature in wt particles (50°C during RT incubation) suggests that the FHV RNA genome 5’ and 3’ proximal regions may be more susceptible to thermodynamic changes, and thus, are prone to be released from capsid shell at lower temperatures; (**B**) eluted and puff particles showed generally overlapping profiles, which entails increased coverage in several isolated regions. This suggests that RNA genome release may prioritize specific regions, instead of the entire genome; (**C**) compared to eluted particles, puff particles only showed substantial increase in coverage at RNA1: 151-250nts, suggesting that RNA1: 151-250nts may be the component of the “puffs” observed in previous CryoEM studies [16].

### Different genomic regions must overcome specific energy barriers prior to release during virion disassembly

To extend PT-ClickSeq beyond the static eluted and puff particle intermediates and to characterize the incremental particle disassembly and genome release, we heated purified FHV virions (ramp rate at +0.1°C/s) to different temperatures (range from 52.5°C to 67.5°C, with 2.5°C interval). After heat-treatment, we again performed PT-ClickSeq and evaluated the read coverage over the viral genome to reveal genomic RNA exposure (**Fig. 3**). As before, read coverage was normalized to the total sequenced depth across the viral genome.

**Figure 3:**
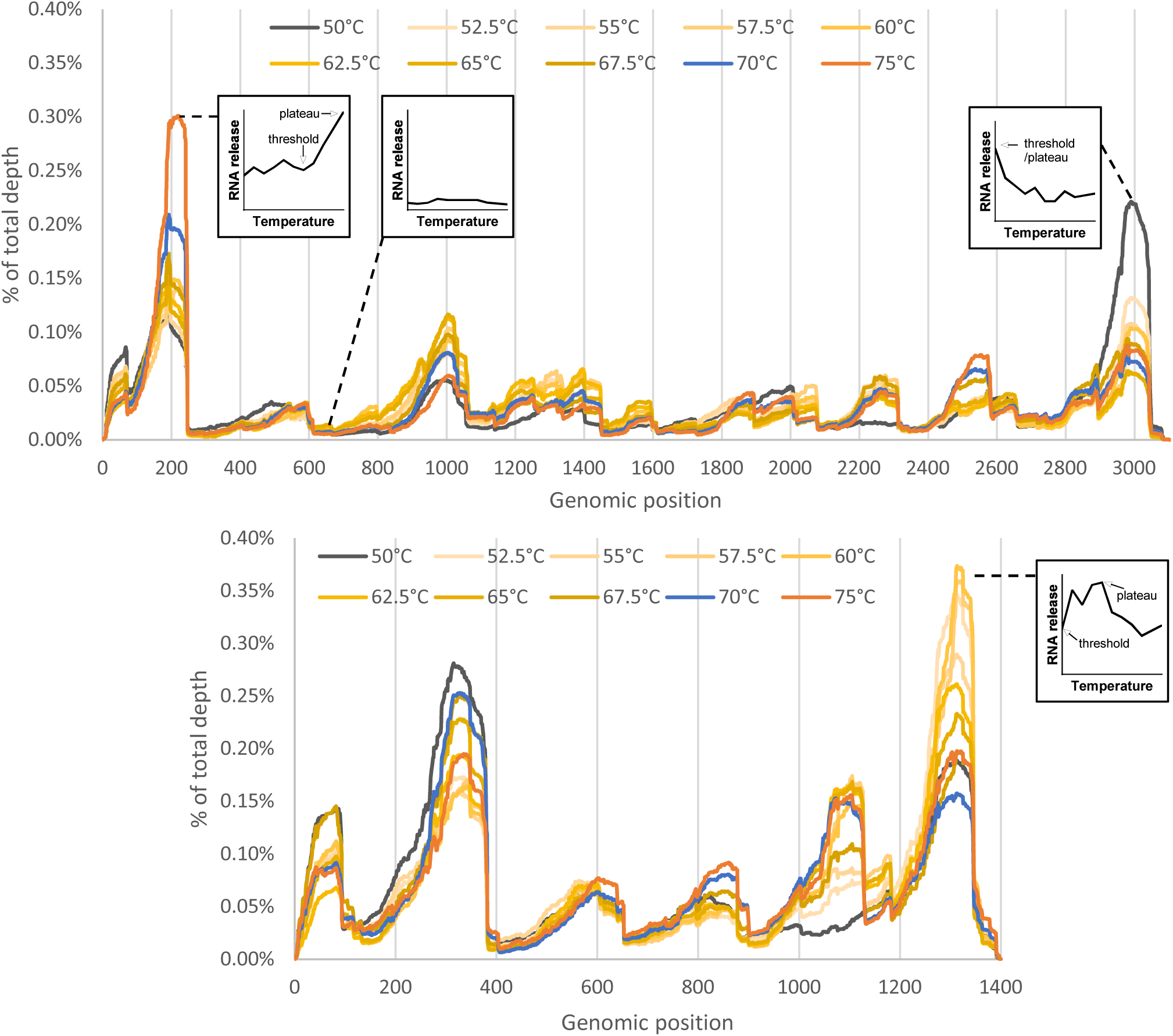
Particle-templated ClickSeq (PT-ClickSeq) reveals release of specific genomic regions at different temperatures. Different FHV genomic regions show differential exposure upon heat treatment at temperatures ranging from 50°C to 75°C. (50°C=wt particle without pre-heating, 70°C=eluted particle, 75°C=puff particle, normalized to % of total depth).

We observed that for both genomic RNAs, different genomic loci showed distinctive trends in read coverage at different temperatures: (**A**) a sequentially increased coverage correlating with temperature (e.g. RNA1: 90-250, RNA2: 1000-1130), indicating continually increased exposure of RNA upon increased temperature treatment; (**B**) generally negative correlation (e.g. RNA1: 1-75, RNA1: 2900-3040), indicating maximal relative coverage early during disassembly and thus having proportionately reduced read coverage upon treatment at higher temperatures; (**C**) unchanged regions of low coverage (e.g. RNA1: 545-700, RNA2: 400-570), indicating consistently protected genomic loci; (**D**) regions of fluctuating coverage during the range of the heat-treatment (e.g. RNA1: 950-1060, RNA2:1-90), suggesting an mid-to-early exposure of viral RNA during an intermediate stage of disassembly. The representative regions, genomic positions and their general trends are presented in **Supplemental Fig. S3**.

This information provides insights to the different energy barriers that must be overcome for each genomic region to be exposed. For example, RNA1: 2150-2310 showed a ∼3.1 fold increase of coverage from 50°C to 52.5°C, which was followed by a gradual fluctuations between 52.5°C and 75°C. This suggests an early to immediate release of this genomic region upon heat-treatment. In contrast, RNA1: 2500-2580 showed minimal changes in read coverage upon heat treatment between 50°C-65°C, but a subsequent increase of read coverage after 67.5°C. The coverage of RNA2: 1250-1350 was relatively unchanged when comparing wt, eluted and puff particles (**Fig.2**). However, upon further investigation, substantial genome release increase can be detected from 50°C to 52.5°C, which remained at a plateau from 52.5°C to 60 °C and then gradually decreased thereafter (**Fig. 3**, **supplemental Fig. S3**). This indicates that the release of RNA2 3’-terminus overcomes the first energy barrier of 52.5°C, and continued to release RNA until genome release peaked at 60°C. The decrease of coverage at temperature treatments >60°C is due to the continued relative increase of other genomic regions during virus particle disassembly.

These data exemplify how different genomic regions, even those separated by less than 200nts of genetic distance, can exhibit different energy barriers prior to their exposure during virus particle disassembly. Altogether, these data indicate that FHV RNA genome release is an ordered and step-wise process, instead of a stochastic or ‘*en masse’* action of release.

### RNA-protein interaction disorganization in FHV disassembly intermediates revealed by vPAR-CL

Viral Photo-Activatable Ribonucleoside CrossLinking (vPAR-CL) is an NGS-based high-throughput method previously developed to study RNA-protein interactions in the context of intact virus particles [20, 29]. Briefly, cells in culture are supplemented with 4-thiouradine (4SU) which is metabolized into 4SUTP and subsequently incorporated into viral genomic RNA during replication. Purified virions with RNA genome(s) containing 4SU can be UV crosslinked to adjacent protein side chains of the viral capsid shell. During RNAseq library generation, the resultant RNA-amino acid adduct is mis-read by the reverse transcriptase as a cytosine. As a result, U-C transitions are found in the final NGS data, revealing the nucleotide position of RNA-protein interactions. Unlike chemical crosslinkers (such as formaldehyde), 4SU/UV induced crosslinking is limited to “zero distance” (does not add an atom) RNA-protein interactions [22], which places vPAR-CL as an ideal method to characterize the RNA-capsid interaction sites in mature virions or during the disassembly process. In addition, vPAR-CL does not rely on the enrichment of crosslinked RNA-protein adduct. Instead, the U-C transition rate is a ratiometric measure of the presence and consistency across a population of conserved RNA-protein interactions. This also places vPAR-CL as a suitable approach to monitor genome-capsid dynamics. As a result, the expected loss of RNA-capsid interaction during FHV disassembly process will be reflected by diminished U-C transition rate across the viral genome or at specific regions, rather than the loss of overall sequence coverage.

As an important validation of the approach, we first assayed the U-C transition rate in the same virus particle samples before (“CL-”) and after (“CL+”) UV crosslinking. (**Fig. 4a**). As expected, wt particles demonstrate higher frequencies of high U-C transition rates in the CL+ samples than CL-controls (confirmed by two-sampled Kolmogorov-Smirnov (K-S) tests, *p* < 2e-16 for RNA1 and *p* < 3e-14 for RNA2). We observed that wt FHV elicited strong and widespread vPAR-CL signals at multiple loci across both RNA1 and RNA2 (**Fig. 4b, Supplementary Fig. S4**). In contrast, the eluted and puff particles both showed diminished overall frequencies of U-C transition rates (K-S tests yielding p-values greater than 0.049).

**Figure 4:**
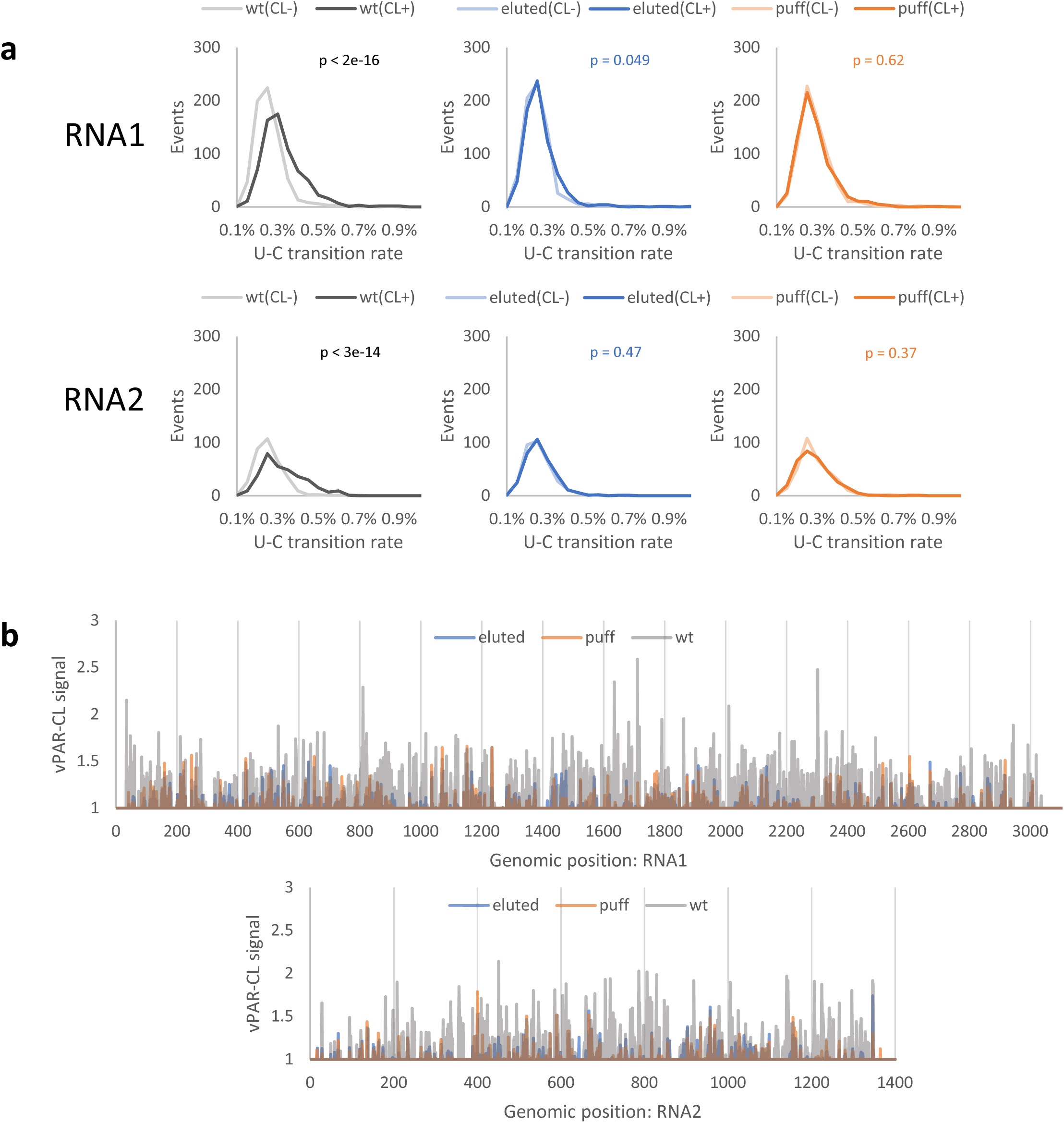
Virus Photoactivatable Ribonucleoside Cross-Linking (vPAR-CL) signals of different disassembly intermediates reveal different RNA-capsid interaction landscapes. (**a**) Amongst different particles, the crosslinked wt particle (“wt(CL+)”) elicited higher U-C transition rate than that of control (“wt(CL-)”, which indicating robust crosslinking at RNA-capsid interaction sites. In contrast, the U-C transition rates of eluted or puff particles were comparable to respective controls. The average U-C transition rates of all U positions were compared between CL+ and CL-, with two sampled Kolmogorov-Smirnov test (N=4). (**b**) The vPAR-CL signals of wt, eluted and puff particles showed overlapping RNA-capsid interaction patterns between eluted and puff particle, but contrasting to the signals of wt particles. Signals generated from the average of 4 independent experiments.

Having established that crosslinking induced specific U-C transitions, we investigated the vPAR-CL signals (positional fold change of U-C transition rate of CL+/CL-) of different particles. Compared to the extensive vPAR-CL signal of wt particles, eluted and puff particles exhibited a clear reduction in vPAR-CL signals across both RNA1 and RNA2 (**Fig. 4b, Supplementary Fig. S4**). This suggests loss of conserved RNA-capsid interactions in eluted and puff particles. Globally, the vPAR-CL signals of eluted and puff particles exhibited a strong linear correlation (Pearson r=0.78) (**Supplementary Fig. S5**). In contract, there was no correlation of vPAR-CL signals between puff and wt particles (Pearson r=0.0047). The overlapping vPAR-CL signals between eluted and puff particles is important, as this suggests that despite the previously observed morphological differences [16], the eluted and puff particles embraced similar internal RNA re-organization and the resultant internal RNA-capsid interactions were consistently retained through morphological transformation and RNA genome release.

Altogether, these data indicate significant re-ordering of encapsidated RNAs and dramatic loss of RNA-capsid interactions in these particles. It is important to note that, although eluted and puff particles exhibited generally reduced vPAR-CL signals compared to wt, several genomic positions retained comparable or even increased vPAR-CL signals (e.g. RNA1:U1233, RNA2:U400). This is consistent with the electrophoretic mobility shift assay (**Fig. 1b**) and suggests that in spite of extensive heating, some virion RNA-capsid interactions were retained.

### FHV genome release during disassembly is anti-corelated with RNA-capsid interactions

The formation of FHV disassembly intermediates demonstrates strong RNA-capsid interactions that survived extensive heating (**Fig. 1**). We sought to understand whether the RNA-capsid interactions in wt FHV contributed to the differential RNA genome release during virus disassembly process (**Fig. 2**). We plotted the vPAR-CL signals of wt particles (>1.5; correspondent to top 15% vPAR-CL signals in wt FHV, **Supplementary Fig. S4**) and % of priming site in wt and puff particles from PT-ClickSeq data (**Fig. 5**). We noticed that the preferentially released RNA sites (indicated by high % of priming site in wt particles) were mostly characterized by a lack of substantial vPAR-CL signals, especially at the immediate downstream of the genome release sites. This is demonstrated in **Supplementary Table S1**, which reports the % of priming sites in different particles and the vPAR-CL signals in wt and puff particles for representative sites in RNA1 and RNA2. We also investigated the correlation between priming rates and vPAR-CL signals. The moving average (15 nts.) of % of priming and vPAR-CL signals in wt particle showed a moderate but significant negative Spearman correlation coefficient (r = −0.35, ANOVA *p* = 1E-90 for RNA1, r = −0.27, ANOVA *p* = 6E-24 for RNA2), suggesting that the RNA genome release at lower temperature (50°C) favored sites without strong RNA-capsid interactions. Similarly, the differentially exposed sites in puff particles also showed significant anti-correlation with vPAR-CL signals (moving average with 15nts window, r = −0.27, ANOVA *p* = 5E-54 for RNA1 and r = −0.31, ANOVA *p* = 1E-31 for RNA2).

**Figure 5:**
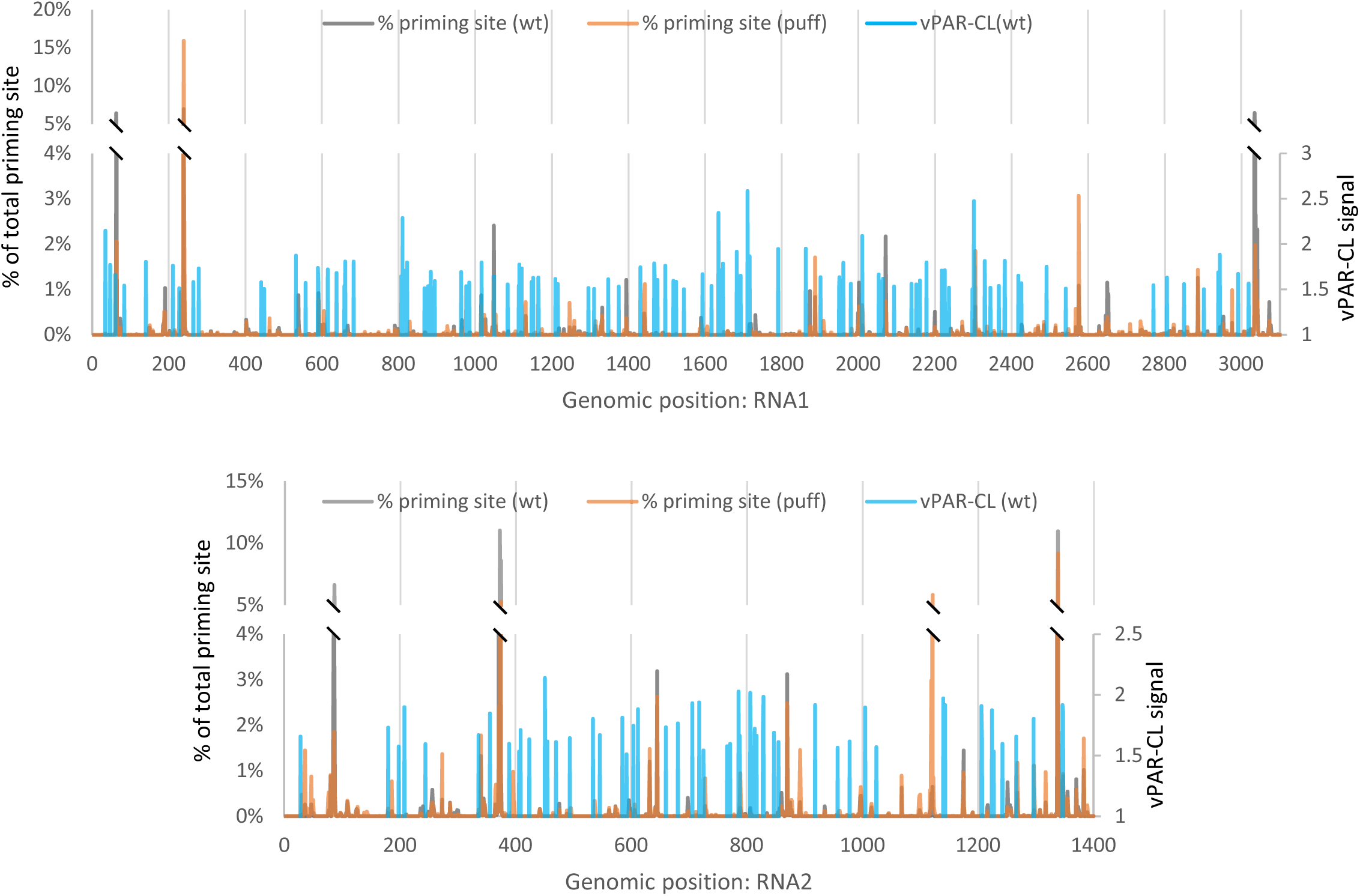
Anti-correlation between exposed genomic regions and RNA-capsid interactions. The exposed genomic regions in wt and puff particles are indicated by the % of total priming sites. These exposed sites are often characterized by a lack of U’s in the sequences immediate downstream, or lack of meaningful RNA-capsid interactions (vPAR-CL signal <1.5).

## Discussion

High-resolution structural studies of intermediate particles of icosahedral virus assembly and disassembly present a significant challenge for advanced imaging technologies such as CryoEM due to their transient nature and conformational heterogeneity. Furthermore, it is often difficult to gain information regarding the encapsidated genetic cargo or the dynamic relationship between RNA and capsid during virus disassembly. Recent studies of virus particles and disassembly intermediates have revealed that the encapsidated viral genome is often asymmetrically ordered within the capsid shell in spite of the imposing structure of the icosahedral capsid shell [16]. In this research, we re-purposed a fragmentation-free RNAseq method “ClickSeq” and demonstrated that it can be used as a convenient tool to study icosahedral virus disassembly intermediates and reveal the sequences of exposed RNA as well as the energetic barriers to release for different genomic regions.

During virus entry and subsequent disassembly, virus particles undergo numerous coordinated and concerted *in vivo* conformational changes that are triggered and dependent upon specific interactions between virus particles and intracellular host factors (e.g. receptor-induced viral uncoating, pH changes in the endosome, etc). These interactions and factors alter the energy landscape of the virus particle in a choreographed manner that results in specific and ordered structural/morphological changes. In this study, FHV disassembly intermediates were obtained by heating wt virions to 70°C or 75°C. Although *in vitro* heat treatment of purified virus particles does not replicate intracellular physiological conditions, such heating of viruses provides a simple and well-controlled method to recapitulate the incremental structural changes that occur during virus particle disassembly under *in vitro* conditions. It has been demonstrated in poliovirus that the conformational changes during heat-induced uncoating fully resemble those *in vivo* after receptor-binding [30]. Heat induced virion structural changes have also been studied among DNA phages [31–33]. In a previous study, a heat-induced FHV RNA-RNA interaction was demonstrated to be important for virus packaging and virion integrity[26].

In this study, using PT-ClickSeq, we revealed that the heat-induced FHV RNA genome release follows a step-wise order and that different genomic regions must overcome different thermodynamic barriers to be exposed (**Fig. 2,3**). During the transition from intact virions to disassembly intermediates (“eluted” and “puff” particles), only specific regions of RNA genome showed significantly increased accessibility. This observation contrasts a possible “*en masse*” model wherein the genetic material is released altogether in a disordered manner. Such a model would result in different genomic loci to be randomly and simultaneously exposed across the virus particles that make up the whole ensemble population, which would yield even read coverage across the viral genome without the features we observed in our study. Further evidence of FHV genome packaging and release can be demonstrated by the vPAR-CL experiments that showed that even under escalated heating conditions, the RNA-capsid interactions are not completely disordered. Instead, strong RNA-capsid interactions survive the disruptive heat and consistent structural tropism was still retained in eluted and puff particles (**Fig. 4, Supplementary Fig. S5**).

From our PT-ClickSeq experiment (**Fig. 2,3**), it is particularly interesting that the 5’ and 3’ termini for both RNA1 and RNA2 are disproportionately exposed at lower temperatures (50°C), where the virus particle is expected to remain fully intact. This observation is reminiscent of the transient exposure, or ‘breathing’ of the membrane-penetrating ‘gamma-peptide’ at ambient temperatures when observed by mass spec and in computational modelling approaches[34, 35]. Similar observations have been demonstrated by a common cold virus (human rhinovirus 2), which is highlighted by the rapidly accessible 3’-end of viral RNA to hybridization methods during heating [36]. These findings suggest a mechanism of icosahedral virus genome egress whereby the free termini of each RNA encounter lower energy barriers to release than the rest of the genome, and therefore can be preferentially exposed during cell entry.

Functionally, it is also conceivable that the rapid accessibility of viral RNA termini plays essential roles in virus replication after cell entry: the 5’-end of +ssRNA viral RNA mimics host mRNAs to recruit the host ribosomal subunits[37, 38]; the 3’-end of +ssRNA virus genome typically serves as initial template for viral genomic or subgenomic RNA replication. Certain characteristics of viral +ssRNA 3’ end (such as polyadenylation) may also function to ensure viral genome stability or to prevent cellular exoribonuclease cleavage [39–41]. For FHV, the of 5’- and 3’-regions of the viral RNA also contain important functional motifs required early during the infection cycles: FHV RNA1 5’-region contains important cis-acting element which directs RNA replication to the mitochondria membrane [42], while the RNA1 3’-end contains the subgenomic RNA3 that encodes silencing suppressor protein B2[43, 44]. The early released RNA1: 2900-3100 region discovered in this study also coincides with FHV 3’ cis elements that are important for the replication of RNA3 and RNA2 [45, 46]. In comparison to RNA1, the motifs of FHV RNA2 are less well characterized. However, the early released RNA2: 1-370 region contains a stem loop structure that is important for RNA2 genome packaging [47] and an RNA1-RNA2 heterodimer stem which are shown to be important for genome packaging specificity as well as overall virion thermostability [19]. RNA2:1200-1400 region also encases a 3’ cis element, responsible for regulating RNA2 replication [48].

Other evidence also suggests that FHV genome release is a gradual, stepwise process instead of abrupt disordering of virus particles. Conformational studies show that FHV genome release may be confined to a single portal at the two-fold axis expansion [16], which echoes the presence of ordered RNA duplexes at the same axis in virus particles [17]. In addition to FHV, the egress of genomic RNA from a single pore (usually located on the symmetrical axis of capsids) on virus particle has also been demonstrated in other icosahedral viruses, such as flaviviruses [49], poliovirus and human enteroviruses [30, 36, 50–53], as well as simulated for virus-like particles [54]. The highly symmetrical nature of non-enveloped virus particles (e.g. an icosahedral particle would encompass 15 identical 2-fold axes) begs the curious question of how the virus designates one portal / axis for genome release. We posit that the asymmetrically ordered sequence(s) and specific RNA-capsid interactions (or the lack-thereof) within a virus particle dictate of the ordered release of genomic material from capsid shell. As such, the encapsidated RNA genome is not passive a cargo, but rather plays an active and ordered role in directing the structural changes required for a virus to successfully infect a host cell.

## Methods

### Cell culture and virus stocks

Wild type (wt) Flock House virus (FHV) was generated by co-transfecting *Drosophila melanogaster* (S2) cells with pMT plasmids containing either FHV RNA1 or RNA2. FHV gene expression was induced with copper sulphate 24 hours after transfection. The transfected cells (P0) were used to inoculate naïve S2 cells to generate continuous passages P1 or P2 of virus stocks. In this study, all FHV stocks used are either P1 or P2 viruses to preserve full-length genomes without accumulation of defective viral genomes [28]. For purification of virus particles, wt FHV stocks were sequentially purified with 4% polyethylene glycol (PEG) 8000 precipitation, RNase/DNase digestion, sucrose gradient ultracentrifugation and 100K MWCO polyether sulfone (PES) membrane filtration. The detailed methods regarding cell culture maintenance, virus passaging, transfection, and purification are previously described [20, 26].

### FHV disassembly intermediates

The heat-induced FHV disassembly intermediates (namely “eluted” and “puff” particles) were generated using previously established methods [16]. Briefly, 50 µg of purified wtFHV particles were resuspended in 50 mM HEPES (pH 7.4) and stored in 4°C. During heating, a thermocycler program was set to heat the wt virions from 4°C to the inducing temperatures (70°C for eluted particle and 75°C for puff particle) at a ramp rate of +0.1°C/s. The particles then were continuously heated at the inducing temperatures for 30 minutes and placed on ice immediately thereafter.

### Particle-templated ClickSeq (PT-ClickSeq)

50 µg of purified particles were used directly as template in reverse transcription reaction, without any RNA extraction. The ClickSeq library preparation generally follows previously developed methods [21, 27], with several modifications to sequenced exposed RNA in native particles. In brief: to preserve the particle integrity, a pre-RT denaturing step was removed (typically at 75°C for 10 mins). Instead, purified particles were mixed with 5 pM of random hexamer primer with a partial i7 Illumina adapter and 1µL of 10 mM AzNTPs:dNTPs mixture. A ratio of 1:5 AzNTPs:dNTPs was used to obtain a short distribution cDNA fragments, which minimizes premature termination of reverse transcription due to possible RNA structure or protein interactions or potential elongation-induced stripping of RNA molecules from remaining capsid shell. The RT reaction was conducted with SuperScript III reverse transcriptase (Invitrogen) with manufacturer protocol and incubated at 50 °C for 40 min. After reverse transcription, the products were treated with RNase H, as per the standard ClickSeq protocol[21]. The 3’-azido-terminated cDNAs were “Click”-ligated to a 5’-alkyne adapter which also contained a 12N unique molecular identifier (UMI). The rest of library preparation, including purification, final PCR amplification, multiplexing was detailed previously [21, 27].

Extracted and purified FHV RNAs were used with the same PT-ClickSeq method and modifications stated above to set a baseline control of read coverage of fully exposed RNA template.

### Bioinformatics

For single-end sequencing data, the raw Illumina reads were first processed with *fastp* [55] to remove Illumina adapters, perform basic quality control, and assign unique molecular identifiers (UMIs) to reads (*-a AGATCGGAAGAGC −l 30 -U –umi_loc read1 –umi_len 14*). This is followed by mapping reads to FHV genome with *bowtie2* [56] (*--local*). In order to control for PCR bias, *umi-tools*[57] was used (dedup -- method=standard). Genome coverage information was analyzed with *samtools* [58] (*mpileup*).

For the pair-end sequencing data, *FASTX_toolkit (*http://hannonlab.cshl.edu/fastx%20toolkit/index.html) was used to apply an additional trimming step to the 5’ of R2 reads, to remove the random hexamer plus one additional nucleotide as a buffer (*fastx_trimmer: -f 8*). The priming site was revealed from the R2 reads using BEDTools [59] (*genomecov -bga −5 -strand +*). Since R2 reads are complementary to R1 or the sense of genome, this reports the 5’-most nucleotide of R2 reads after trimming (the 5’-8^th^ nucleotide before trimming), which also corresponds to the 3’-most nucleotide on genomic sense.

### vPAR-CL

Passage 1 (P1) FHV was used to infect naïve cells at an MOI=1. As previously described [20], 100 µM of 4-thiouridine (Sigma-Aldrich, in DMSO) was supplied to infected cells at both 0 hpi and 16hpi, to reach a final concentration of 200 µM. Infected cells and supernatant were harvested at 40 hpi. To fully release intracellular virions, cell/supernatant was supplemented with 1% Triton X-100 and was allowed to go though one freeze-thaw cycle. 4SU-containing FHV particles were purified with the method stated above.

The purified 4SU-containing FHV (4SU+) was then heated to induce disassembly intermediates with the method described above. After heating and generating of disassembly intermediates, each type of particles was further purified with 100K MWCO (PES) membrane filtration, which removes any free capsid/RNA fragments to prevent non-specific crosslinking. For each particle type (wt, eluted, puff), the 4SU-containing virus was separated into two pools: one pool underwent UV crosslinking (365 nm at 0.15 J/cm^2^), proteinase K digestion (8U, 37°C for 30min) and RNA extraction (Zymo Direct-zol RNA kit) to yield the crosslinked particles (CL+); and the other pool only underwent proteinase K digestion and RNA extraction to yield the non-crosslinked controls (CL-).

The NGS library preparation of RNA samples extracted from CL+ and CL-particles are described previously [20]. One important improvement is that this study utilized a 12N UMI adapter to be click-ligated with azido-terminated cDNA, which allows for de-duplicating and bioinformatic control PCR artifacts. Raw sequencing reads were processed the same way as single-end reads as stated above. Pileup files were generated using *samtools*, and the nucleotide frequency and mismatch rate at each genomic position were analyzed with *vPARCL-call* (https://github.com/andrewrouth/vPARCL_call).

The vPAR-CL signal is defined as the fold change between crosslinked samples (CL+) and non-crosslinked controls (CL-). Biological triplicates of both CL+ and CL-were included for each particle type. In order to ensure the quality of vPAR-CL data, the minimum sequencing depth threshold was set to 1000 de-duplicated reads.

## Data Availability

The raw FASTQ sequencing data of this study are available in the NCBI sequence read archive (SRA) with accession number: PRJNA1098227.

---

**Supplemental Figure S1: Differential genome coverage of wt, eluted and puff particles**. (**a**) The summed percentage of coverage of specific genomic regions. (**b**) Each datapoint represents the summed percentage of coverage of a 50nts window (e.g. nt1-50).

**Supplemental Figure S2: Rationale of using paired-end sequencing to reveal priming sites**. Single-end sequencing reveals information from the 5’ of the ‘forward’ read of stranded libraries (R1). The priming site in the reverse-transcription reaction is located at the 5’-end of the ‘reverse’ read of stranded libraries (R2).

**Supplemental Figure S3: Representative regions with different temperature sensitivity.** This table shows the representative regions of Figure 4, their representative genomic positions (in brackets) and general trends of coverage at the representative positions. Red dot: peak coverage.

**Supplemental Figure S4: Virus Photoactivatable Ribonucleoside Cross-Linking (vPAR-CL) signals of wt, eluted or puff particles**. Signals generated from the average of 4 independent experiments. The average coverage showed even sequencing depth and comparable read coverage between control groups (CL-) and crosslinked groups (CL+).

**Supplemental Figure S5: eluted and puff particles showed comparable RNA-capsid interaction patterns**. (a) vPAR-CL signals of puff and eluted particles showed substantial overlap. (b) vPAR-CL signals of eluted and puff particles showed strong linear correlation (Pearson r=0.78). This is contrasted to the non-linear correlation between the vPAR-CL profiles of puff and wt particles (Pearson r = −0.06).

## References

1. Suomalainen, M. and U.F. Greber, Uncoating of non-enveloped viruses. Curr Opin Virol, 2013. 3(1): p. 27–33.

2. Hogle, J.M., Poliovirus cell entry: common structural themes in viral cell entry pathways. Annu Rev Microbiol, 2002. 56: p. 677–702.

3. Brandenburg, B., et al., Imaging poliovirus entry in live cells. PLoS Biol, 2007. 5(7): p. e183.

4. Tsang, S.K., et al., Kinetic analysis of the effect of poliovirus receptor on viral uncoating: the receptor as a catalyst. J Virol, 2001. 75(11): p. 4984–9.

5. Fuchs, R. and D. Blaas, Uncoating of human rhinoviruses. Rev Med Virol, 2010. 20(5): p. 281–97.

6. Wetz, K. and T. Kucinski, Influence of different ionic and pH environments on structural alterations of poliovirus and their possible relation to virus uncoating. J Gen Virol, 1991. 72 (Pt 10): p. 2541–4.

7. Brabec, M., et al., Conformational changes, plasma membrane penetration, and infection by human rhinovirus type 2: role of receptors and low pH. J Virol, 2003. 77(9): p. 5370–7.

8. Walukiewicz, H.E., J.E. Johnson, and A. Schneemann, Morphological Changes in the T=3 Capsid of Flock House Virus during Cell Entry. Journal of Virology, 2006. 80(2): p. 615.

9. Odegard, A.L., et al., Low endocytic pH and capsid protein autocleavage are critical components of Flock House virus cell entry. J Virol, 2009. 83(17): p. 8628–37.

10. Greber, U.F. and M. Way, A superhighway to virus infection. Cell, 2006. 124(4): p. 741–54.

11. Burckhardt, C.J. and U.F. Greber, Virus movements on the plasma membrane support infection and transmission between cells. PLoS Pathog, 2009. 5(11): p. e1000621.

12. Agirre, J., et al., Cryo-electron microscopy reconstructions of triatoma virus particles: a clue to unravel genome delivery and capsid disassembly. J Gen Virol, 2013. 94(Pt 5): p. 1058–1068.

13. Tuthill, T.J., et al., Equine rhinitis A virus and its low pH empty particle: clues towards an aphthovirus entry mechanism? PLoS Pathog, 2009. 5(10): p. e1000620.

14. Dormitzer, P.R., et al., Structural rearrangements in the membrane penetration protein of a non-enveloped virus. Nature, 2004. 430(7003): p. 1053–8.

15. Chandran, K., et al., The delta region of outer-capsid protein micro 1 undergoes conformational change and release from reovirus particles during cell entry. J Virol, 2003. 77(24): p. 13361–75.

16. Azad, K. and M. Banerjee, Structural dynamics of non-enveloped virus disassembly intermediates. J Virol, 2019.

17. Fisher, A.J. and J.E. Johnson, Ordered duplex RNA controls capsid architecture in an icosahedral animal virus. Nature, 1993. 361(6408): p. 176–9.

18. Odegard, A., M. Banerjee, and J.E. Johnson, Flock house virus: a model system for understanding non-enveloped virus entry and membrane penetration. Curr Top Microbiol Immunol, 2010. 343: p. 1–22.

19. Zhou, Y. and A.L. Routh, Bipartite viral RNA genome heterodimerization influences genome packaging and virion thermostability. J Virol, 2024. 98(3): p. e0182023.

20. Zhou, Y. and A. Routh, Mapping RNA-capsid interactions and RNA secondary structure within virus particles using next-generation sequencing. Nucleic Acids Res, 2020. 48(2): p. e12.

21. Routh, A., et al., ClickSeq: Fragmentation-Free Next-Generation Sequencing via Click Ligation of Adaptors to Stochastically Terminated 3’-Azido cDNAs. Journal of molecular biology, 2015. 427(16): p. 2610–2616.

22. Zhou, Y., S.L. Sotcheff, and A.L. Routh, Next-generation sequencing: A new avenue to understand viral RNA-protein interactions. J Biol Chem, 2022. 298(5): p. 101924.

23. Schwarcz, W.D., et al., Virus stability and protein-nucleic acid interaction as studied by high-pressure effects on nodaviruses. Cell Mol Biol (Noisy-le-grand), 2004. 50(4): p. 419–27.

24. Schneemann, A., V. Reddy, and J.E. Johnson, The Structure and Function of Nodavirus Particles: A Paradigm for Understanding Chemical Biology. 1998. 50: p. 381–446.

25. Tihova, M., et al., Nodavirus Coat Protein Imposes Dodecahedral RNA Structure Independent of Nucleotide Sequence and Length. Journal of Virology, 2004. 78(6): p. 2897–2905.

26. Zhou, Y. and A.L. Routh, Bipartite viral RNA genome heterodimerization influences genome packaging and virion thermostability. bioRxiv, 2022: p. 2022.07.29.501896.

27. Jaworski, E. and A. Routh, ClickSeq: Replacing Fragmentation and Enzymatic Ligation with Click-Chemistry to Prevent Sequence Chimeras, in Next Generation Sequencing: Methods and Protocols, S.R. Head, P. Ordoukhanian, and D.R. Salomon, Editors. 2018, Springer New York: New York, NY. p. 71–85.

28. Jaworski, E. and A. Routh, Parallel ClickSeq and Nanopore sequencing elucidates the rapid evolution of defective-interfering RNAs in Flock House virus. PLoS Pathog, 2017. 13(5): p. e1006365.

29. Sung, P.Y., et al., A multidisciplinary approach to the identification of the protein-RNA connectome in double-stranded RNA virus capsids. Nucleic Acids Res, 2023.

30. Bostina, M., et al., Poliovirus RNA is released from the capsid near a twofold symmetry axis. J Virol, 2011. 85(2): p. 776–83.

31. Duda, R.L., et al., Structure and energetics of encapsidated DNA in bacteriophage HK97 studied by scanning calorimetry and cryo-electron microscopy. J Mol Biol, 2009. 391(2): p. 471–83.

32. Kawai, Y., Thermal transition profiles of bacteriophage T4 and its DNA. J Gen Appl Microbiol, 1999. 45(3): p. 135–138.

33. Qiu, X., Heat Induced Capsid Disassembly and DNA Release of Bacteriophage λ. PLOS ONE, 2012. 7(7): p. e39793.

34. Bothner, B., et al., Evidence of viral capsid dynamics using limited proteolysis and mass spectrometry. J Biol Chem, 1998. 273(2): p. 673–6.

35. Jana, A.K. and E.R. May, Atomistic dynamics of a viral infection process: Release of membrane lytic peptides from a non-enveloped virus. Sci Adv, 2021. 7(16).

36. Harutyunyan, S., et al., Viral uncoating is directional: exit of the genomic RNA in a common cold virus starts with the poly-(A) tail at the 3’-end. PLoS Pathog, 2013. 9(4): p. e1003270.

37. Jaafar, Z.A. and J.S. Kieft, Viral RNA structure-based strategies to manipulate translation. Nat Rev Microbiol, 2019. 17(2): p. 110–123.

38. Decroly, E., et al., Conventional and unconventional mechanisms for capping viral mRNA. Nat Rev Microbiol, 2011. 10(1): p. 51–65.

39. Kim, D., et al., The Architecture of SARS-CoV-2 Transcriptome. Cell, 2020. 181(4): p. 914–921 e10.

40. Barr, J.N. and R. Fearns, How RNA viruses maintain their genome integrity. Journal of General Virology, 2010. 91(6): p. 1373–1387.

41. Dreher, T.W., Functions of the 3ʹ-untranslated regions of positive strand RNA viral genomes. Annual review of phytopathology, 1999. 37(1): p. 151–174.

42. Van Wynsberghe, P.M. and P. Ahlquist, 5’ cis elements direct nodavirus RNA1 recruitment to mitochondrial sites of replication complex formation. J Virol, 2009. 83(7): p. 2976–88.

43. Li, H., W.X. Li, and S.W. Ding, Induction and suppression of RNA silencing by an animal virus. Science, 2002. 296(5571): p. 1319–21.

44. Eckerle, L.D., C.G. Albarino, and L.A. Ball, Flock House virus subgenomic RNA3 is replicated and its replication correlates with transactivation of RNA2. Virology, 2003. 317(1): p. 95–108.

45. Lindenbach, B.D., J.Y. Sgro, and P. Ahlquist, Long-distance base pairing in flock house virus RNA1 regulates subgenomic RNA3 synthesis and RNA2 replication. J Virol, 2002. 76(8): p. 3905–19.

46. Miller, D.J., et al., Engineered Retargeting of Viral RNA Replication Complexes to an Alternative Intracellular Membrane. J Virol, 2003. 77(22): p. 12193–12202.

47. Zhong, W.D., R. Dasgupta, and R. Rueckert, Evidence That the Packaging Signal for Nodaviral Rna2 Is a Bulged Stem Loop. Proceedings of the National Academy of Sciences of the United States of America, 1992. 89(23): p. 11146–11150.

48. Albariño, C.G., L.D. Eckerle, and L.A. Ball, The cis-acting replication signal at the 3ʹ end of Flock House virus RNA2 is RNA3-dependent. Virology, 2003. 311(1): p. 181–191.

49. Skubnik, K., et al., Capsid opening enables genome release of iflaviruses. Sci Adv, 2021. 7(1).

50. Levy, H.C., et al., Catching a virus in the act of RNA release: a novel poliovirus uncoating intermediate characterized by cryo-electron microscopy. J Virol, 2010. 84(9): p. 4426–41.

51. Shingler, K.L., et al., The enterovirus 71 A-particle forms a gateway to allow genome release: a cryoEM study of picornavirus uncoating. PLoS Pathog, 2013. 9(3): p. e1003240.

52. Belnap, D.M., et al., Molecular tectonic model of virus structural transitions: the putative cell entry states of poliovirus. J Virol, 2000. 74(3): p. 1342–54.

53. Hadfield, A.T., et al., The refined structure of human rhinovirus 16 at 2.15 Å resolution: implications for the viral life cycle. Structure, 1997. 5(3): p. 427–441.

54. Sukenik, L., et al., Cargo Release from Nonenveloped Viruses and Virus-like Nanoparticles: Capsid Rupture or Pore Formation. ACS Nano, 2021. 15(12): p. 19233–19243.

55. Chen, S., et al., fastp: an ultra-fast all-in-one FASTQ preprocessor. Bioinformatics, 2018. 34(17): p. i884–i890.

56. Langmead, B., et al., Ultrafast and memory-efficient alignment of short DNA sequences to the human genome. Genome biology, 2009. 10(3): p. R25–R25.

57. Smith, T., A. Heger, and I. Sudbery, UMI-tools: modeling sequencing errors in Unique Molecular Identifiers to improve quantification accuracy. Genome Res, 2017. 27(3): p. 491–499.

58. Li, H., et al., The Sequence Alignment/Map format and SAMtools. Bioinformatics (Oxford, England), 2009. 25(16): p. 2078–2079.

59. Quinlan, A.R. and I.M. Hall, BEDTools: a flexible suite of utilities for comparing genomic features. Bioinformatics, 2010. 26(6): p. 841–2.

