## Supplemental Figures for "Ordered release of genomic RNA during icosahedral virus disassembly"

#### - Supplementary Figures

Yiyang Zhou[1,2,*], Andrew L. Routh[2,3,4,5,*]

1. Department of Microbiology and Immunology, The University of Texas Medical Branch, Galveston, Texas, USA
2. Department of Biochemistry and Molecular Biology, The University of Texas Medical Branch, Galveston, Texas, USA
3. Department of Immunology and Microbiology, Scripps Research, La Jolla, California, USA
4. Sealy Center for Structural Biology and Molecular Biophysics, The University of Texas Medical Branch, Galveston, Texas, USA
5. Institute for Human Infections and Immunity, University of Texas Medical Branch, Galveston, Texas, USA

*To whom correspondence should be addressed.

#### Supplemental Figure S1

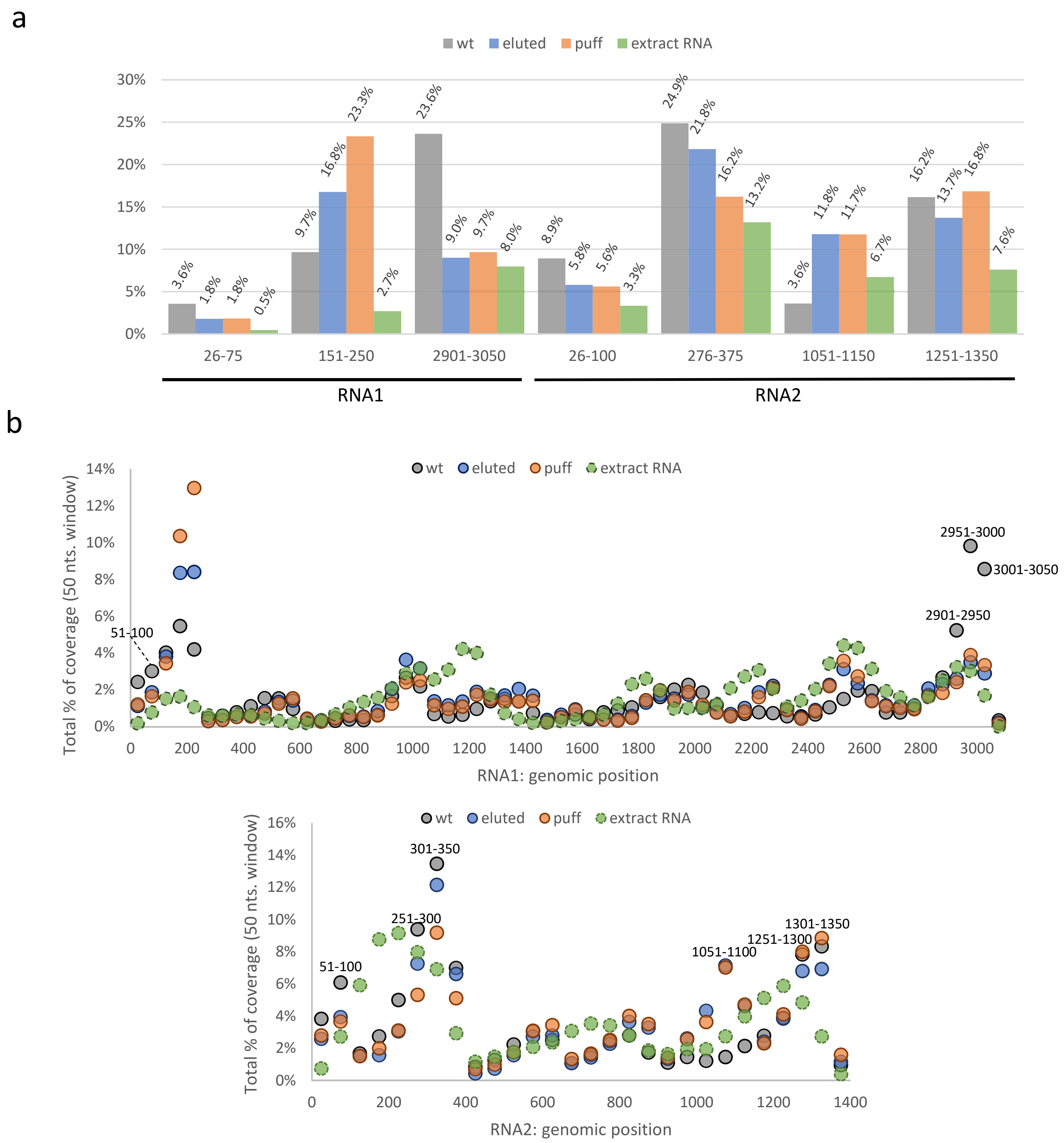


**Supplemental Figure S1: Differential genome coverage of wt, eluted and puff particles**. (**a**) The summed percentage of coverage of specific genomic regions. (**b**) Each datapoint represents the summed percentage of coverage of a 50nts window (e.g. nt1-50).

#### Supplemental Figure S2

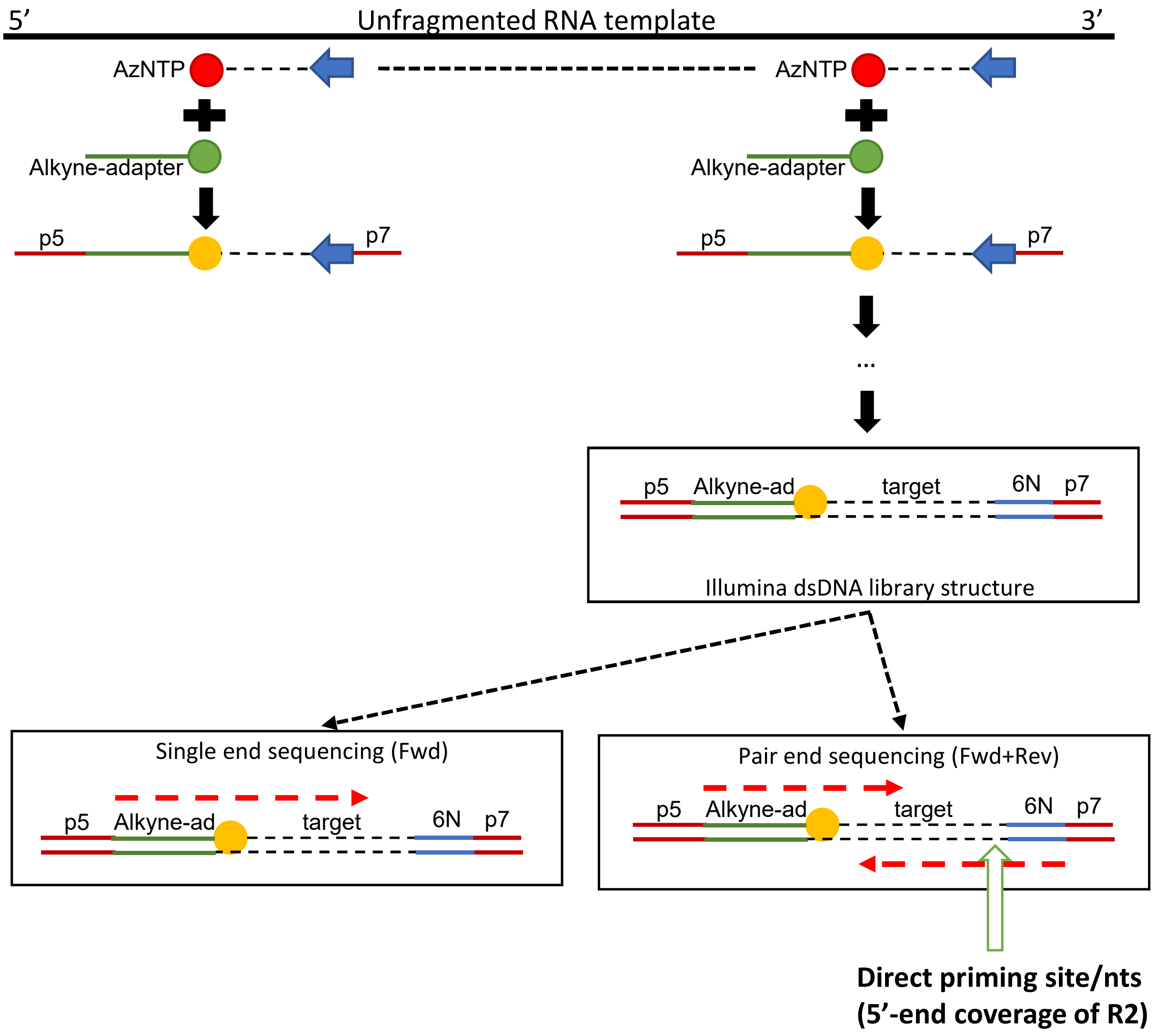


**Supplemental Figure S2: Rationale of using paired-end sequencing to reveal priming sites**. Single-end sequencing reveals information from the 5' of the 'forward' read of stranded libraries (R1). The priming site in the reverse-transcription reaction is located at the 5'-end of the 'reverse' read of stranded libraries (R2).

### Supplemental Figure S3.

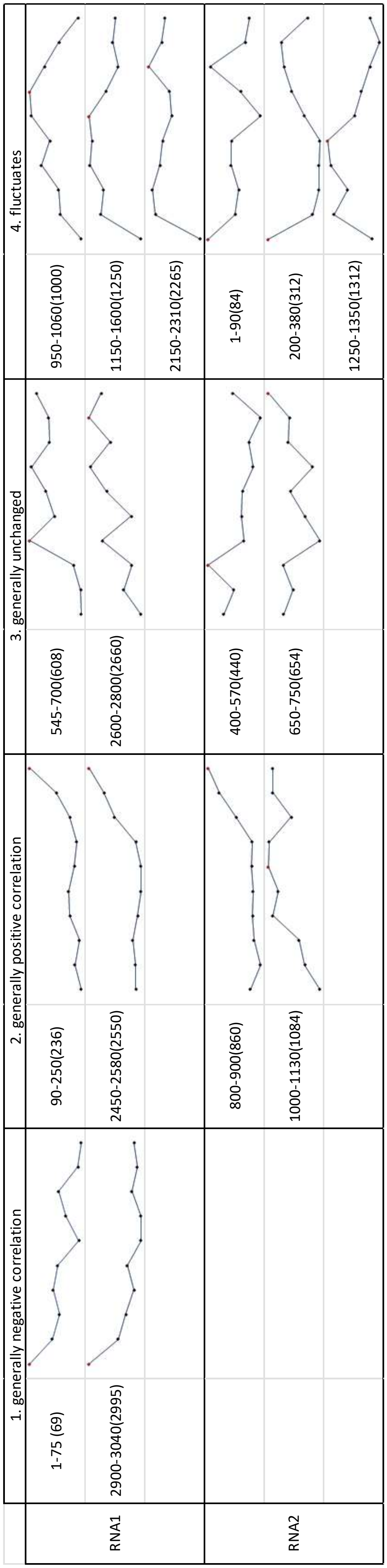


**Supplemental Figure S3: Representative regions with different temperature sensitivity.** This table shows the representative regions of **Figure 4**, their representative genomic positions (in brackets) and general trends of coverage at the representative positions. Red dot: peak coverage.

#### Supplemental Figure S4

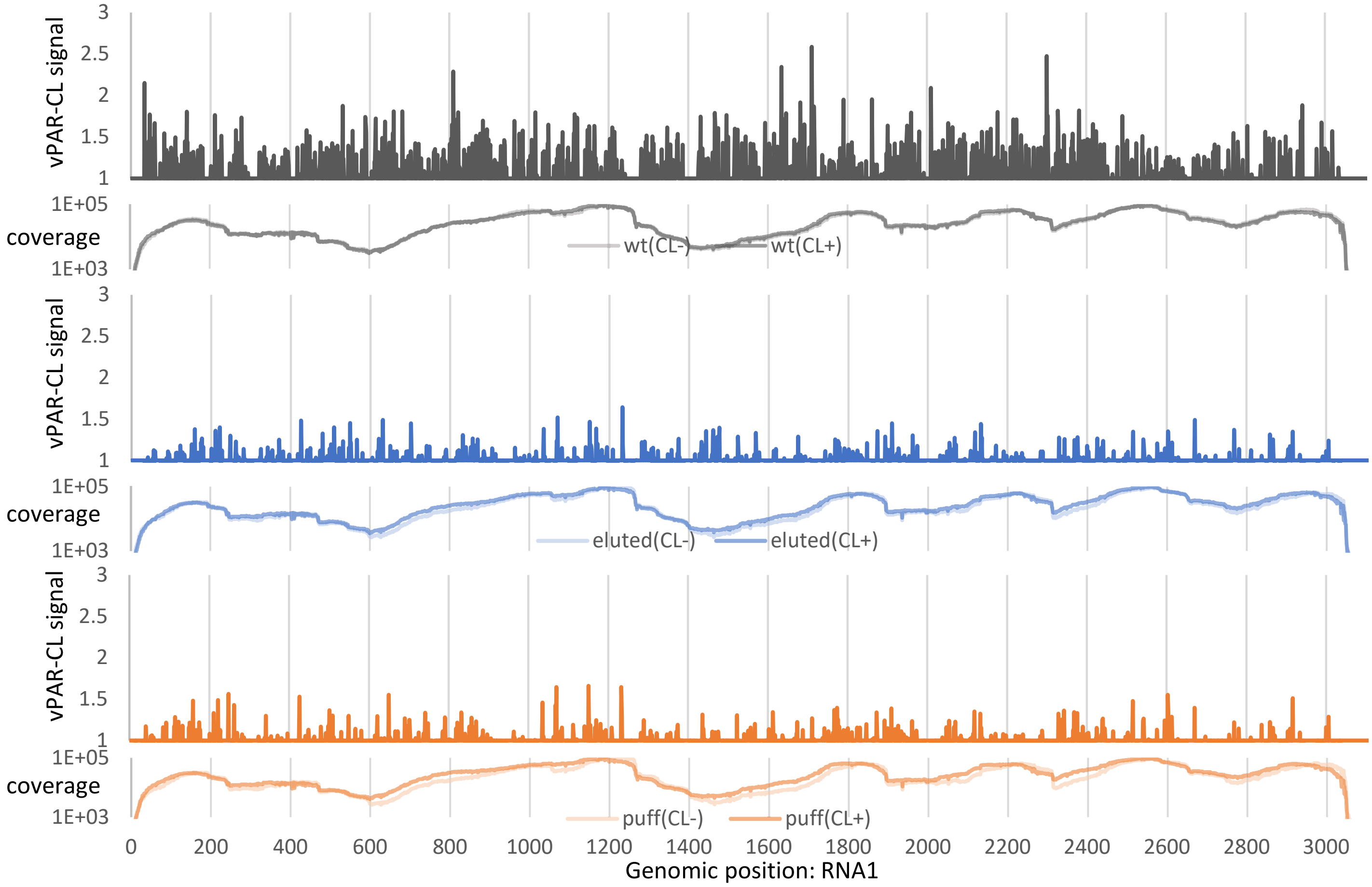


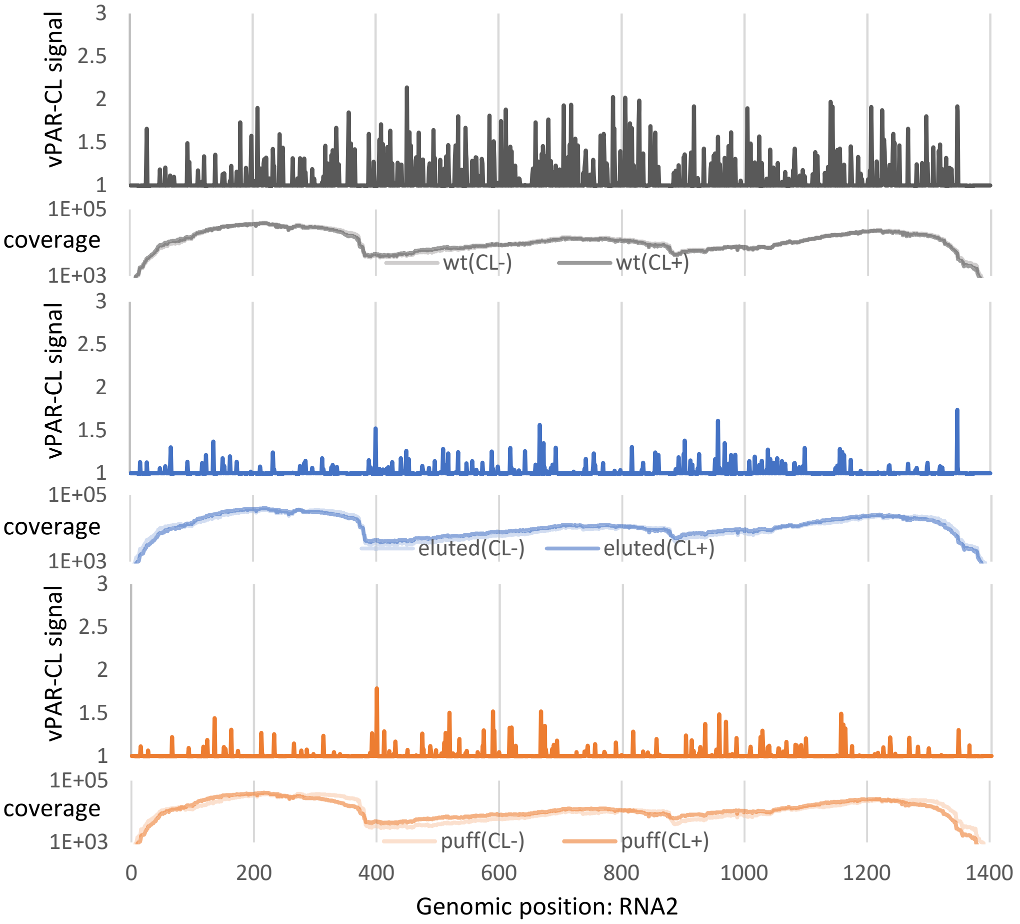

**Supplemental Figure S4: Virus Photoactivatable Ribonucleoside Cross-Linking (vPAR-CL) signals of wt, eluted or puff particles**. Signals generated from the average of 4 independent experiments. The average coverage showed even sequencing depth and comparable read coverage between control groups (CL-) and crosslinked groups (CL+).

#### Supplemental Figure S5

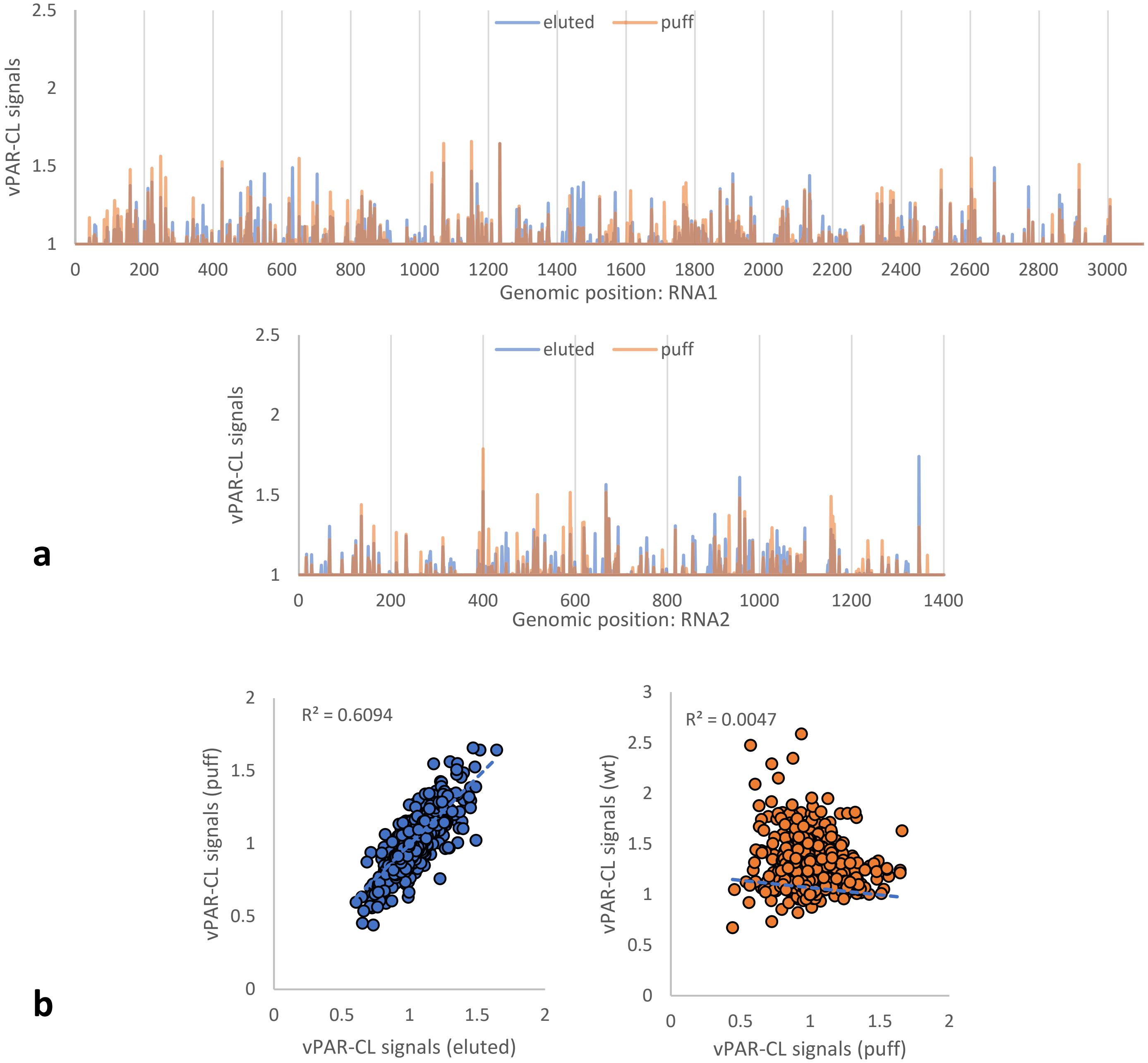


**Supplemental Figure S5: eluted and puff particles showed comparable RNA-capsid interaction patterns**. (a) vPAR-CL signals of puff and eluted particles showed substantial overlap. (b) vPAR-CL signals of eluted and puff particles showed strong linear correlation (Pearson r=0.78). This is contrasted to the non-linear correlation between the vPAR-CL profiles of puff and wt particles (Pearson r = -0.06).
